## Supplementary Information for "Molecular insights into substrate translocation in an elevator-type metal transporter"

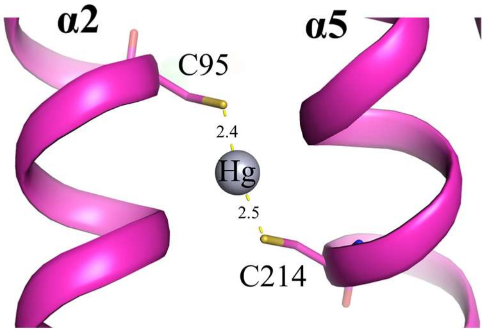


**Figure S1.** $Hg^{2+}$-mediated cross-linking of the A95C/A214C variant. The distance between the indicated atoms are shown in angstrom.

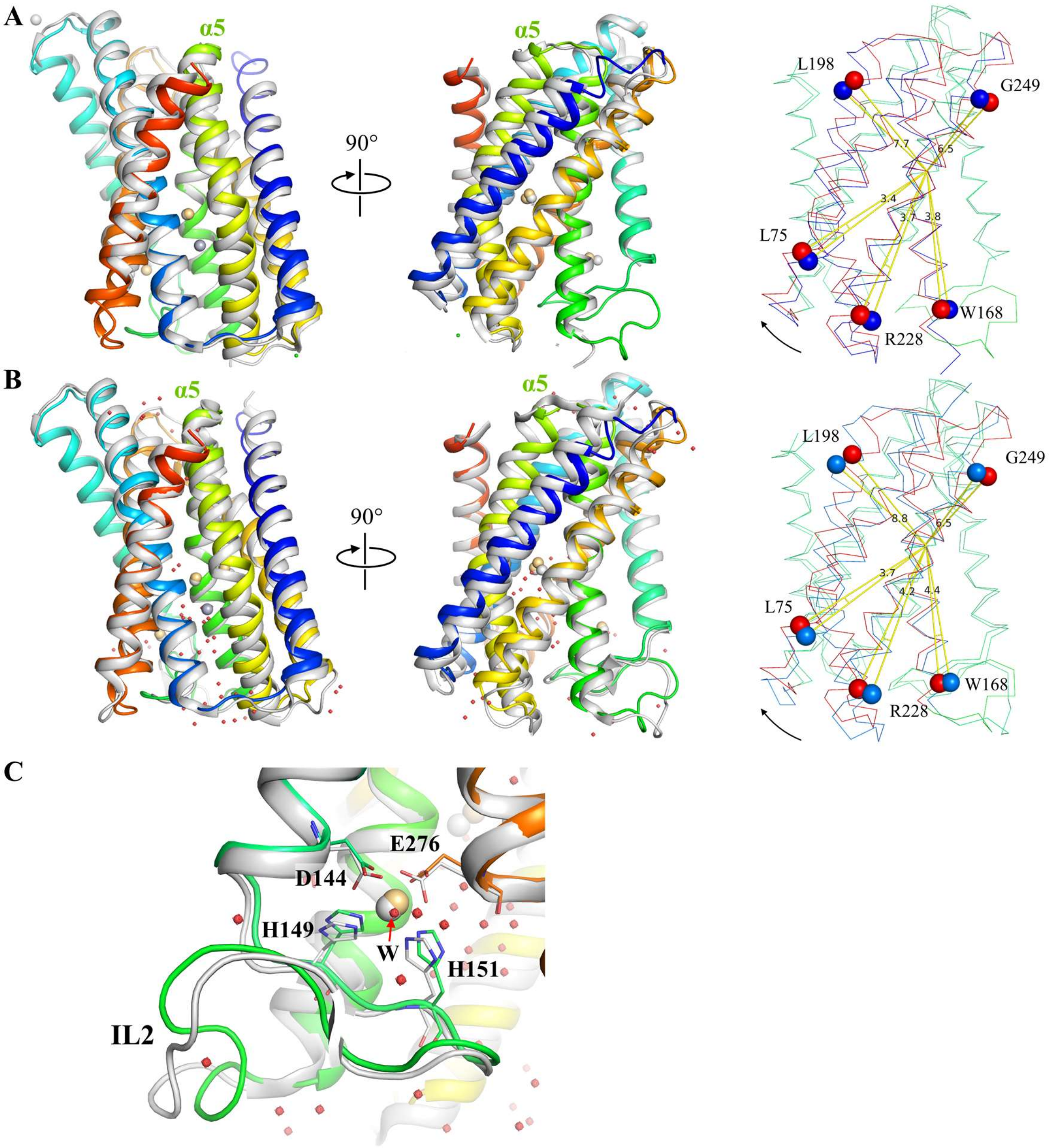


**Figure S2.** Structural comparison of the $Hg^{2+}$-cross-linked A95C/A214C variant (colored in rainbow) with other representative IFCs (in white). (**A**) Structural superimposition with the crystal structure of wild-type BbZIP in the Cd-bound state (PDB: 5TSB) with an R.M.S.D. of 0.96 Å. (**B**) Structural superimposition with the cryo-EM structure of wild-type BbZIP in the Cd-bound state (PDB: 8GHT, chain A) with an R.M.S.D. of 1.00 Å. To align the structures in (A) and (B), the scaffold domains (TM 2/3/7/8) were superimposed to reveal a rotation of the transport domain (TM 1/4/5/6) of the cross-linked structure in which TM5 (yellow-green) undergoes the greatest upward shift (up to 3 Å) than the other TMs. The very N-terminal TM0 is trimmed for clarity. The right panels in (A) and (B) show a quantitative analysis of the upward hinge motion of the transport domain relative to the scaffold domain when the cross-linked structure (red) is compared to other IFC structures (blue). The rotation angels (in degree) at five positions are labeled. The hinge motion is approximately centered at P180. (**C**) Comparison of the M3 sites in the cross-linked structure (rainbow) and the cryo-EM structure (grey). Some differences are noted, including monodentate (cross-linked protein) vs. bidentate coordination (cryo-EM) for D144, different orientations of the carboxylate groups of D144 and E276, and a coordinated water in the structure of the cross-linked protein. The conformations of IL2 (with a Cα RMSD of 1.76 Å) are more different than the other parts of the structure, indicating greater flexibility. The metal chelating residues are shown in stick mode. Water molecules revealed in the cross-linked structure are depicted as small red spheres and the red arrow indicates the coordinated water.

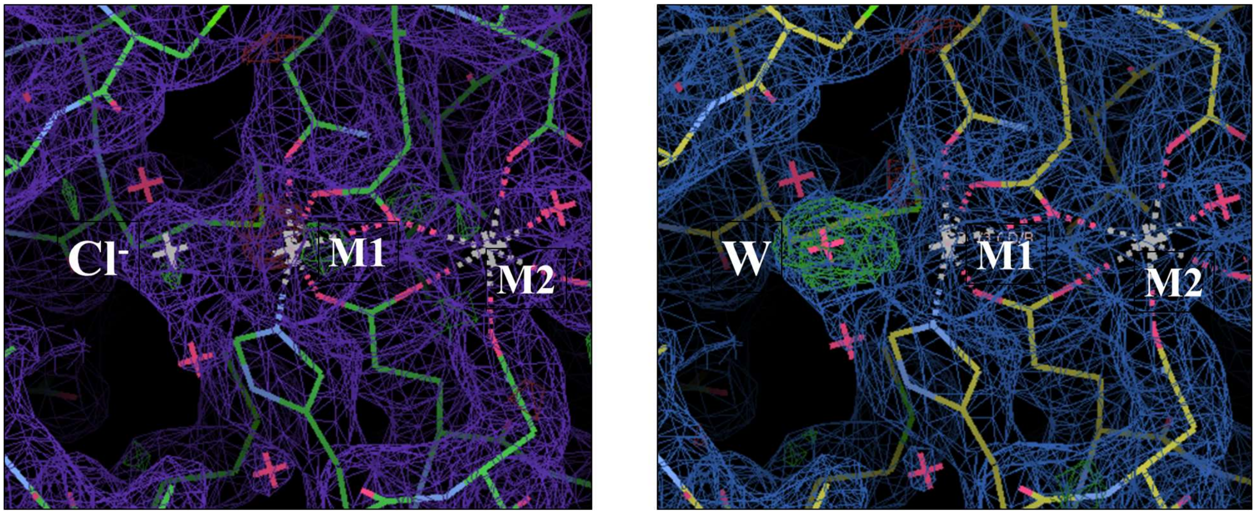


**Figure S3.** Electron density of the chloride ion coordinating the $Cd^{2+}$ bound at the M1 site. *Left*: the density map with a modeled chloride ion ($Cl^-$). *Right*: density map with a modeled water molecule (W). Blue/purple meshes: 2Fo-Fc map at σ=1; Green meshes: positive densities in the Fo-Fc map at σ=3. An ordered water molecule was excluded because of the otherwise residual positive density as indicated in the Fo-Fc difference map. After examining possible ligands present in the crystallization solution containing 300 mM $Cl^-$, only $Cl^-$ matches the density with a *B* factor of 43 comparable to those of the surrounding atoms (30-50) and the distance between the density and $Cd^{2+}$ (2.6 Å).

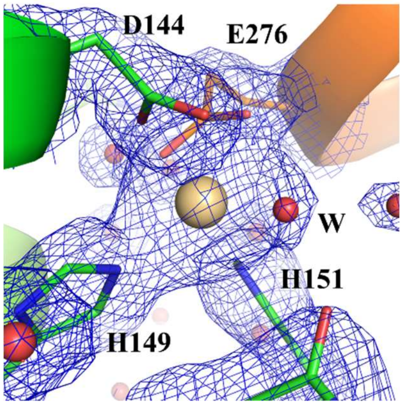


**Figure S4.** Zoom-in view of the electron density at the M3 site (2Fo-Fc map, σ=4). Waters are depicted as red spheres. The bound $Cd^{2+}$ was found to coordinate with a water molecule (W), in addition to four residues (D144, E276, H149, and H151).

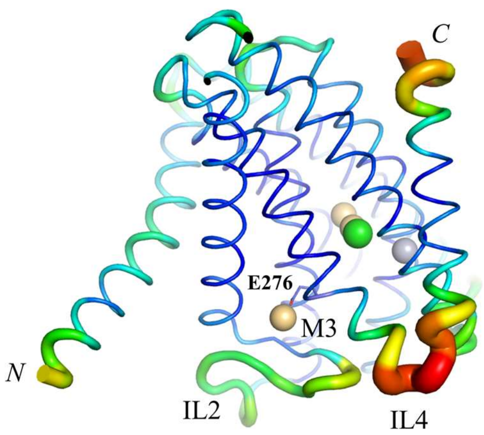


**Figure S5.** B-factor putty of the $Hg^{2+}$-cross-linked structure of the A95C/A214C variant. IL2 and IL4 show great level of flexibility. E276 is shown in stick mode. $Cd^{2+}$, $Hg^{2+}$, and $Cl^-$ are depicted as spheres in light brown, grey, and green, respectively.

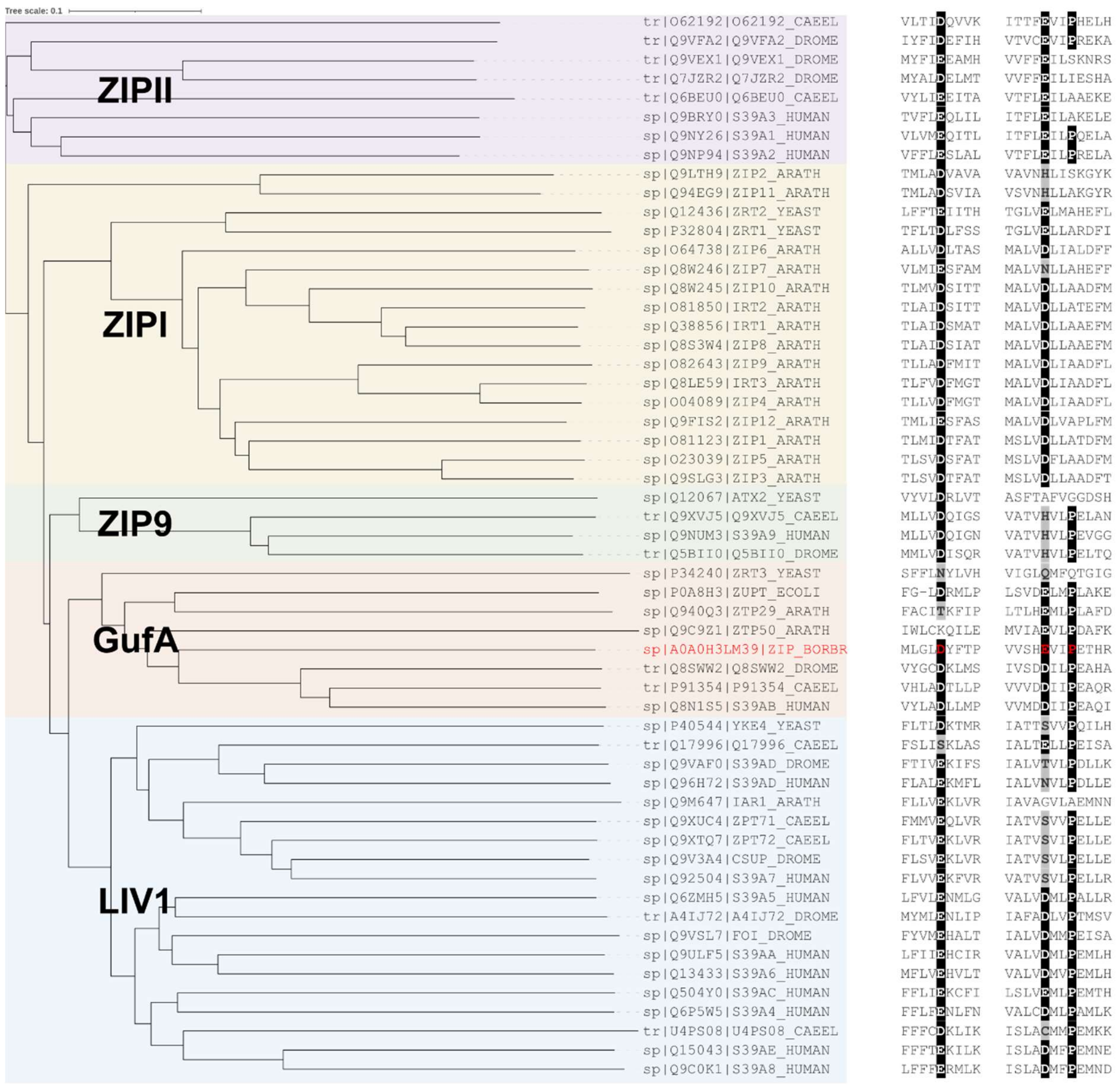


**Figure S6.** Sequence alignment of ZIPs from representative species. A total of 56 ZIPs, including 14 from *Homo sapiens*, 9 from *Drosophila melanogaster*, 5 from *Saccharomyces cerevisiae*, 8 from *Caenorhabditis elegans*, 18 from *Arabidopsis thaliana*, and two bacterial ZIPs, were aligned using Clustal Omega (https://www.ebi.ac.uk/jdispatcher/msa/clustalo). The resulting unrooted phylogenetic tree was visualized at iTOL (https://itol.embl.de/). Major subfamilies are marked with different colors. Only the segments containing the residues topologically equivalent to D144, E276, and P279 in BbZIP (in red) are shown. For the residues corresponding to D144 or E276, glutamates and aspartates are shown in white on a black background and other residues potentially involved in metal binding are on a grey background. Only the P279-equivalent proline residues are highlighted.

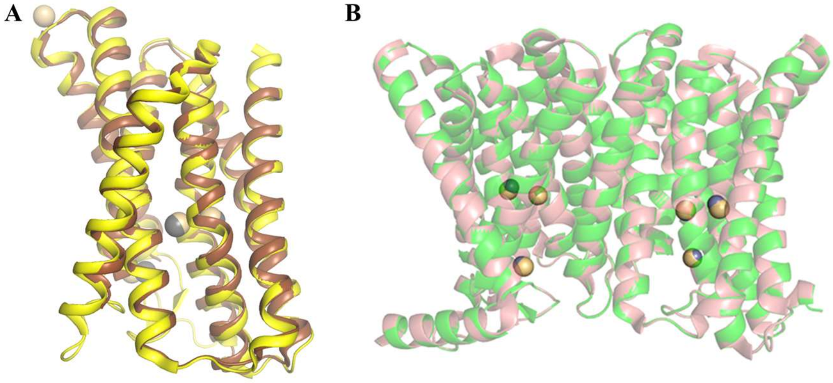


**Figure S7.** Comparison of the relevant structures. (**A**) Superimposition of the crystal structure of the Cd-bound BbZIP (PDB: 5TSB, brown) and the cryo-EM structure of the Cd-bound BbZIP (PDB: 8GHT, yellow). $Cd^{2+}$ are shown as spheres in light brown (5TSB) or grey (8GHT). The Cα RMSD of two structures is 0.68 Å. (**B**) Superimposition of the structure of the Cd-bound BbZIP (PDB: 8GHT, pink) and the model of the Zn-bound BbZIP for simulations (green). $Cd^{2+}$ and $Zn^{2+}$ are depicted as yellow and gray spheres, respectively.

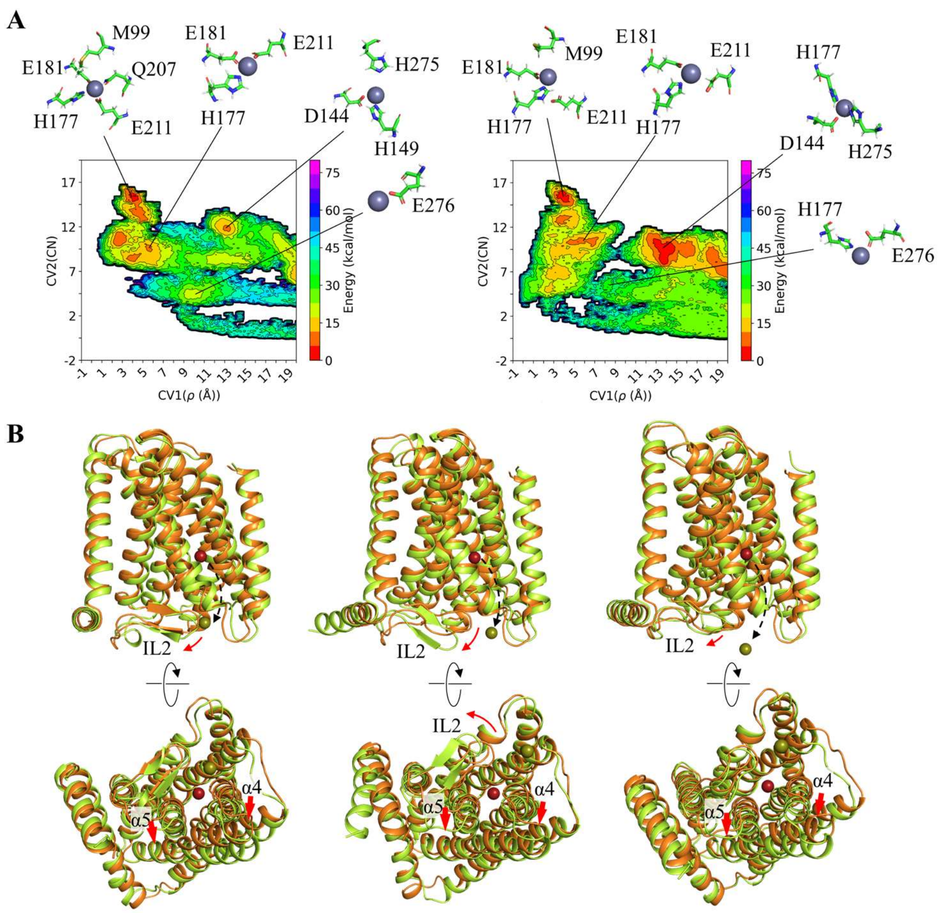


**Figure S8.** Metal release into the cytoplasm in MTD simulations. (**A**) Free energy surface maps of metal coordination during the simulations in the first scenario. The results of one simulation are shown in Figure 4A, and the results of the additional two simulations are shown here. CV1 represents the distance from the metal ion to the protein center of mass, and CV2 is the number of non-hydrogen atoms within 4 Å of the metal. (**B**) Structural rearrangements (red arrows) of the IL2 and the cytoplasmic portions of TM4 and TM5 upon metal released into the cytoplasm via Path 2 (dashed curve arrow) in the second scenario. The structural modes before and after metal release into the cytoplasm are colored in orange and green, respectively. The results of one simulation are shown in Figure 4C, and the results of the additional three simulations are shown here.

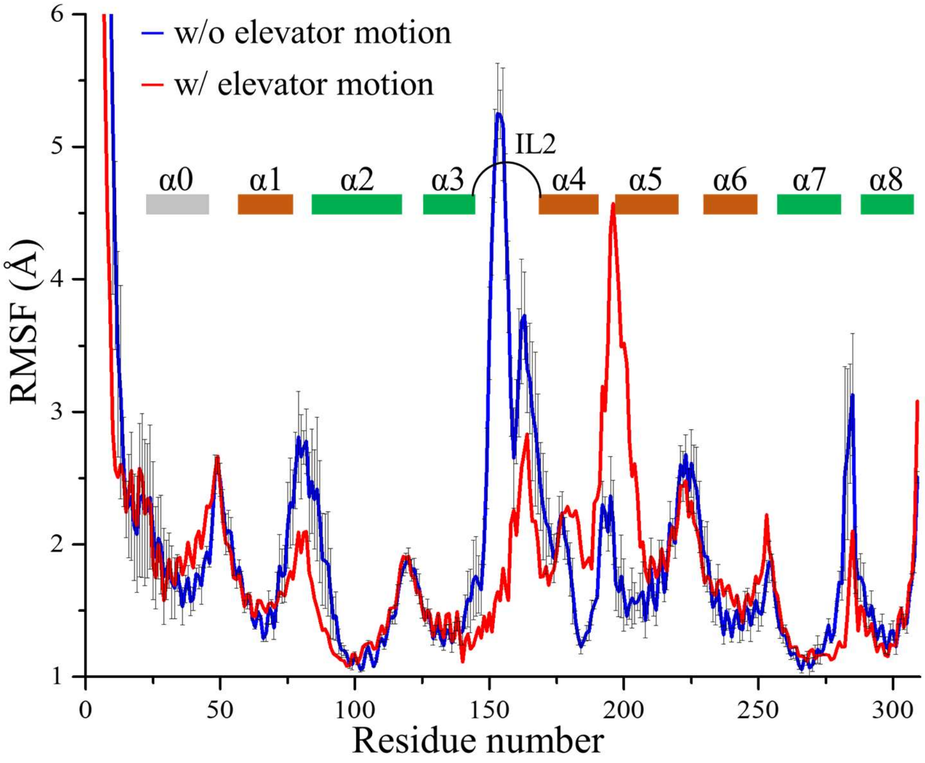


**Figure S9.** RMSF analysis of the simulations with and without elevator motion. The RMSF of each residue is plotted against the residue number. The blue profile represents the mean of three independent simulations in the first scenario where no elevator motion is observed. The black error bars represent the standard errors. The red profile represents the RMSF of the simulation in which the elevator motion was observed in the second scenario. The TMs are indicated by rectangles with the transport domain in green and the scaffold domain in brown.

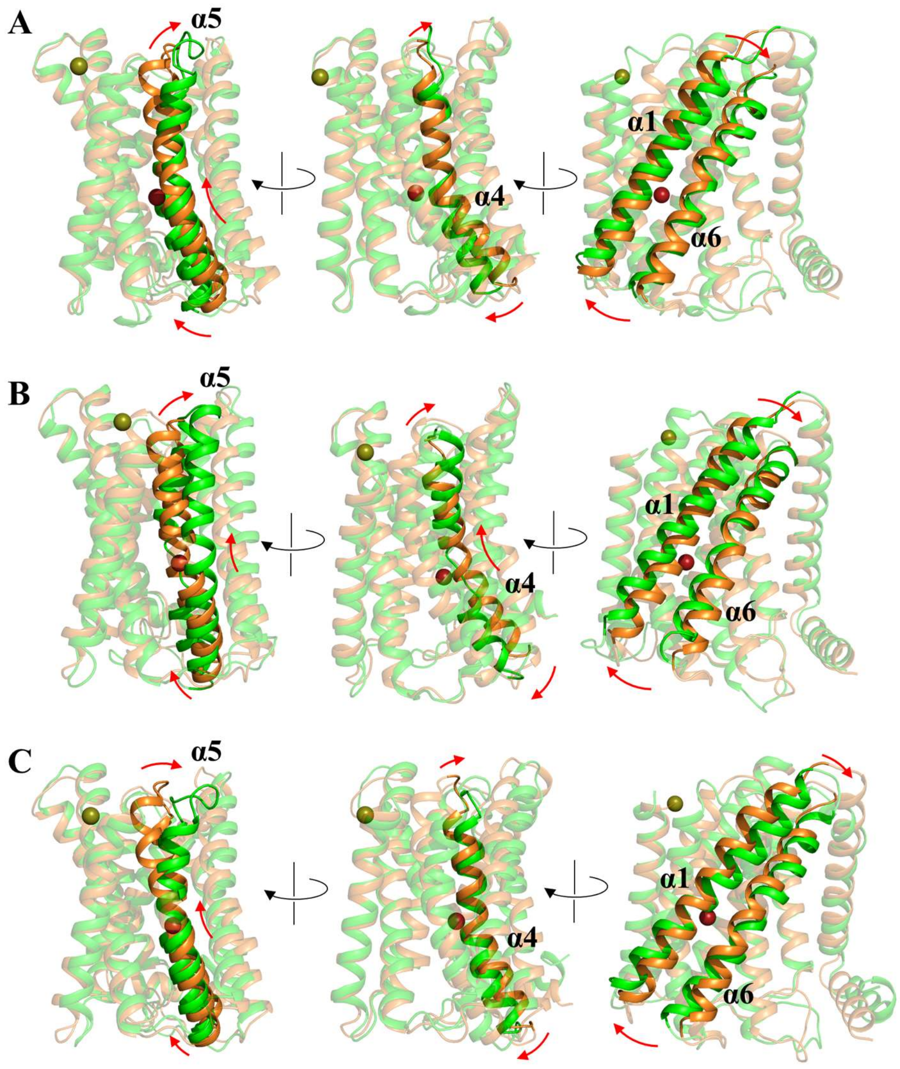


**Figure S10.** Elevator motion of the transport domain observed in MTD simulations in the third (**A**) and the last (**B**, **C**) scenarios. The conformational changes of the highlighted structural elements are indicated by red arrows. The dark red and yellow-green spheres represent $Zn^{2+}$ before and after the IFC-to-OFC transition, respectively.

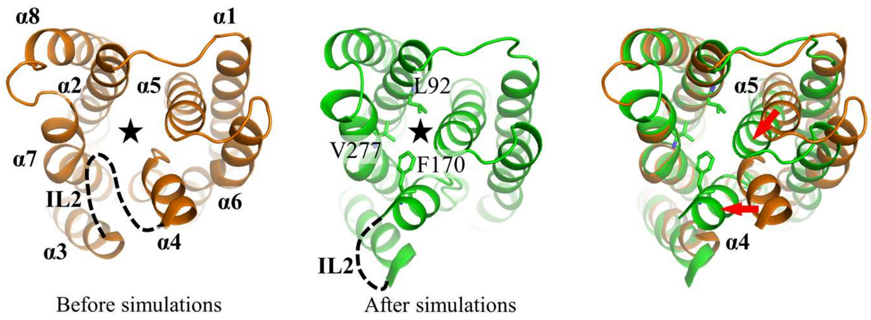


**Figure S11.** Closing of the cytoplasmic gate by the portions of TM4 and TM5 on the cytoplasmic side after removal of the IL2 in standard MD simulations. The hydrophobic residues at the cytoplasmic gate (black star) are labeled.

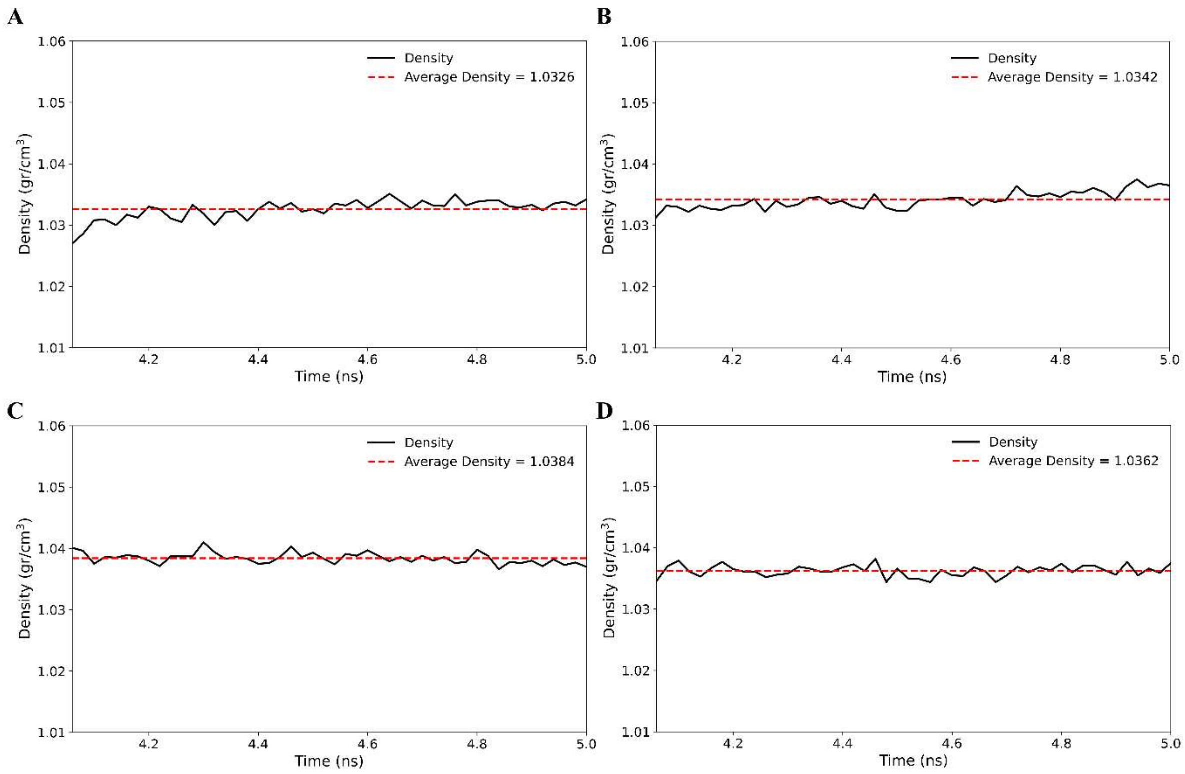


**Figure S12.** Density of each system during the final 1 ns of the equilibration phase. Panels A, B, C, and D represent the scenarios from first to last, respectively.

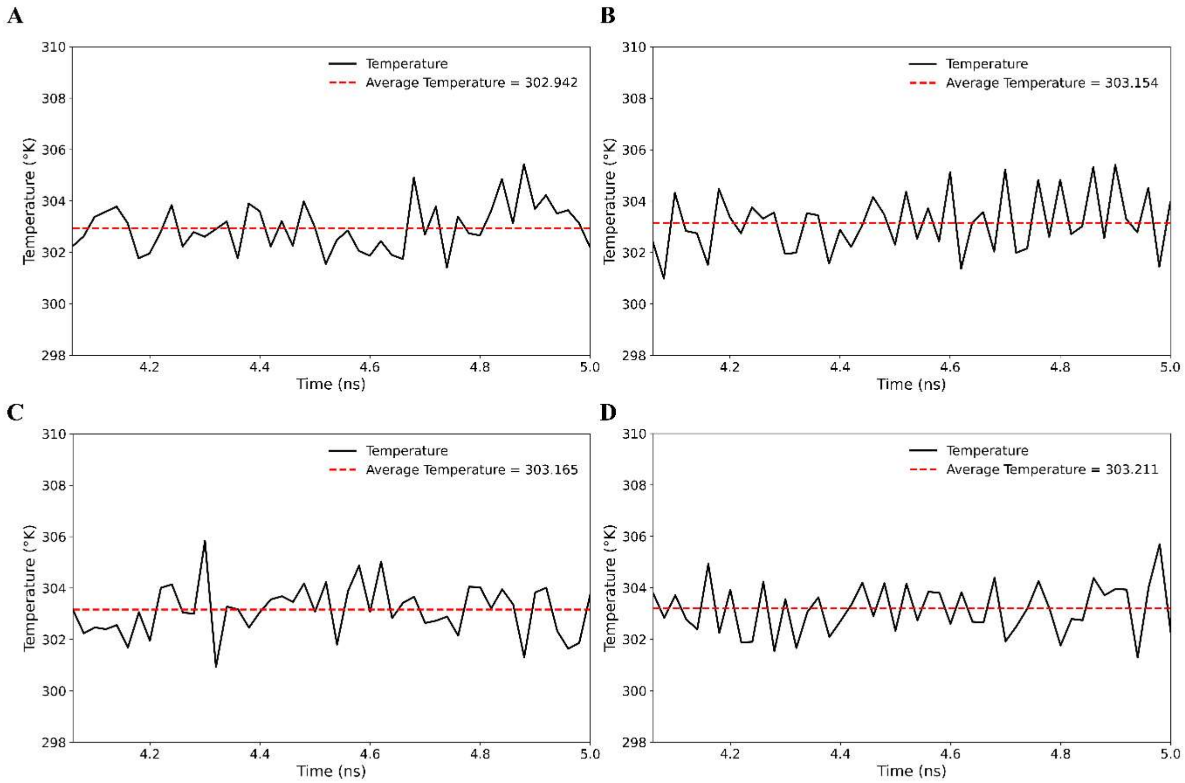


**Figure S13.** Temperature of each system during the final 1 ns of the equilibration phase. Panels A, B, C, and D represent the scenarios from first to last, respectively.

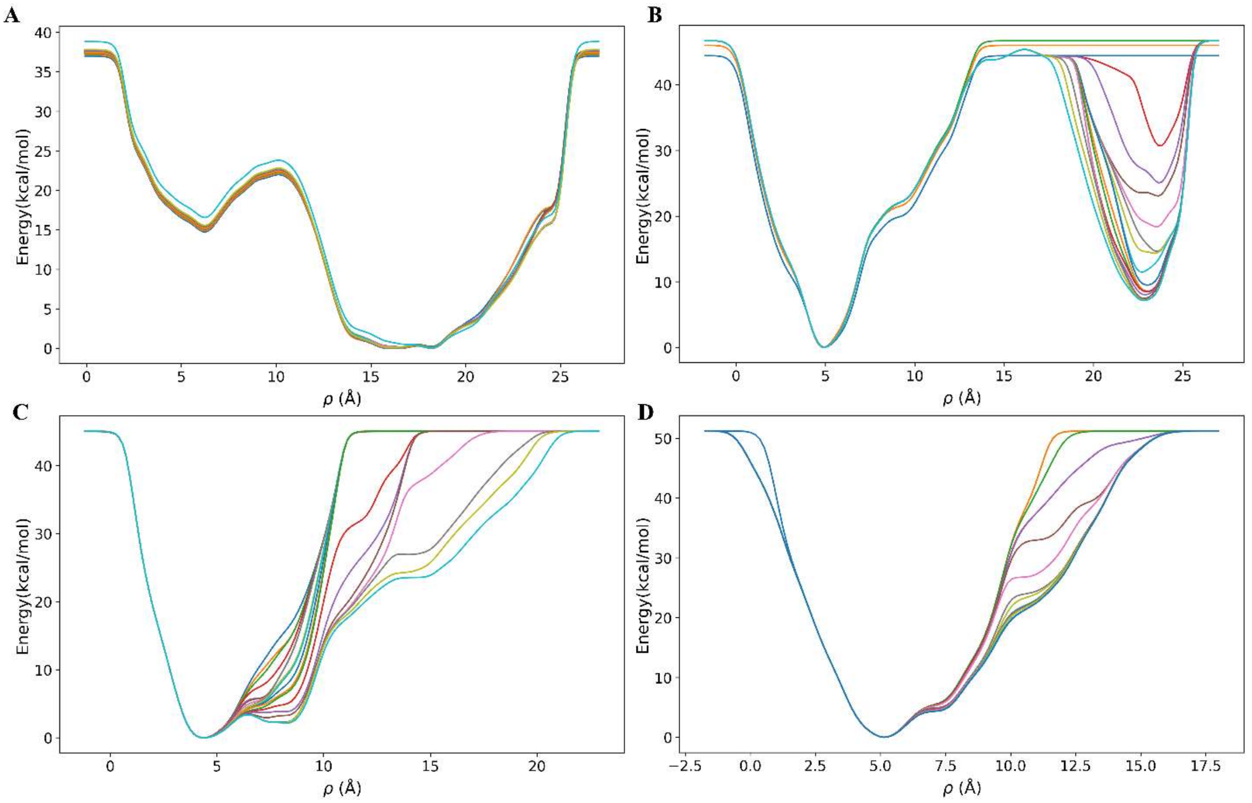


**Figure S14.** Convergence of each system in the MTD simulations. Panels A, B, C, and D correspond to the first, second, third, and fourth scenarios, respectively. The plots are generated every 10 ns during the last 200 ns of the MTD simulations. In the present case, the convergence analysis should primarily focus on the region of real space within the metal release tunnel of the transporter, which corresponds to a CV rho value of approximately 15 Å (at the level of the M3 site). In this region, the bias deposition remains nearly constant over the last ~200 ns indicating that sufficient sampling has occurred in the relevant region. Beyond this the ion proceeds into solution resulting in expected variability in the plots.

**Table S1.** Crystallographic statistics

| **Crystal** | $Hg^{2+}$-crosslinked A95C/A214C BbZIP |
|---|---|
| **Data collection** | |
| Beamline | GM/CA-CAT (23-ID-B) |
| Wavelength (Å) | 0.987 |
| Space group | C $2_1$ |
| Unit cell | |
| a, b, c (Å) | 116.4, 51.0, 51.8 |
| α, β, γ (°) | 90, 113.9, 90 |
| [a]Resolution (Å) | 30.4 - 1.95 (2.02 – 1.95) |
| [a]Redundancy | 10.3 (2.3) |
| [a]Completeness (%) | 97.0 (78.4) |
| [a]$I/\sigma I$ | 14.3 (0.9) |
| [a,b]$R_{merge}$ | 0.172 (0.769) |
| [a,c]$R_{pim}$ | 0.05 (0.475) |
| [d]$CC_{1/2}$ of the highest resolution shell | 0.566 |
| **Refinement** | |
| Unique reflections | 19668 |
| Number of Atoms | 2217 |
| Protein | 2028 |
| Ligands | 131 |
| $H_2O$ | 58 |
| [e]$R_{work}/R_{free}$ | 0.1712/0.1999 |
| *Wilson B-factor* ($Å^2$) | 40.71 |
| *B*-factors ($Å^2$) | |
| Protein | 39.32 |
| Ligands | 59.94 |
| $H_2O$ | 45.9 |
| R.m.s. deviations | |
| Bond lengths (Å) | 0.007 |
| Bond angles (°) | 0.94 |
| Ramachandran plot (%) | |
| Favored | 97.91 |
| Allowed | 2.06 |
| Outliers | 0.0 |

[a]Highest resolution shell is shown in parentheses.

[b]$R_{merge} = \sum_{hkl} \sum_j |I_j(hkl) - \langle I(hkl)\rangle| / \sum_{hkl} \sum_j I_j(hkl)$, where $I$ is the intensity of reflection.

[c]$R_{pim} = \sum_{hkl} [1/(N-1)]^{1/2} \sum_j |I_j(hkl) - \langle I(hkl)\rangle| / \sum_{hkl} \sum_j I_j(hkl)$, where N is the redundancy of the dataset.

[d]$CC_{1/2}$ is the correlation coefficient of the half datasets.

[e]$R_{work} = \sum_{hkl} | |F_{obs}| - |F_{calc}| | / \sum_{hkl} |F_{obs}|$, where $F_{obs}$ and $F_{calc}$ is the observed and the calculated structure factor, respectively. $R_{free}$ is the cross-validation R factor for the test set of reflections (5% of the total) omitted in model refinement.

**Table S2.** Summary of the results of the MTD simulations conducted in this work.

| Conditions | Simulation replicates | Simulation length (ns) | Elevator motion to OFC | Metal release or retention |
|---|---|---|---|---|
| M1 and M2 loaded with $Zn^{2+}$; M3 empty; IL2 unfolded (First scenario) | 1 | 600 | No | Cytoplasm |
| | 2 | 500 | No | Cytoplasm |
| | 3 | 500 | No | Cytoplasm |
| M1, M2, and M3 loaded with $Zn^{2+}$; IL2 folded (Second scenario) | 1 | 500 | No | Cytoplasm* |
| | 2 | 500 | No | Cytoplasm* |
| | 3 | 500 | **Yes** | **Periplasm** |
| | 4 | 500 | No | Cytoplasm* |
| | 5 | 500 | No | Cytoplasm* |
| M1 and M2 loaded with $Zn^{2+}$; M3 empty; IL2 folded (Third scenario) | 1 | 500 | No | Remains around M1 |
| | 2 | 500 | No | Cytoplasm* |
| | 3 | 500 | **Yes** | **Periplasm** |
| M1 and M3 loaded with $Zn^{2+}$; M2 empty; IL2 folded (Last scenario) | 1 | 700 | No | Remains at M1 |
| | 2 | 500 | No | Moves to and remains at M2 |
| | 3 | 500 | No | Cytoplasm* |
| | 4 | 500 | **Yes** | **Periplasm** |
| | 5 | 700 | **Yes** | **Periplasm** |

*Accompanied with conformational changes of the IL2 and the portions of TM4 and TM5 on the cytoplasmic side.

**Table S3.** Detailed information about each MTD simulation system.

| Conditions | # of water molecules | Salt concentration (M) | Lipid composition (POPE:POPG, molar ratio) | Box dimension (x*y*z) (Å) | # of atoms |
|---|---|---|---|---|---|
| First scenario | 18827 | 0.15 | 3:1 | 90* 89*115 | 90510 |
| Second scenario | 16346 | 0.15 | 3:1 | 92*89*104 | 83052 |
| Third scenario | 16351 | 0.15 | 3:1 | 90* 89*105 | 83069 |
| Last scenario | 15916 | 0.15 | 3:1 | 90*90*103 | 81761 |
